# Pro-longevity compounds extend *Caenorhabditis elegans* male lifespan and reproductive healthspan

**DOI:** 10.1101/2025.06.29.662228

**Authors:** Rose S. Al-Saadi, Patrick C. Phillips

## Abstract

Sex differences in aging are robust and ubiquitous. Demographic differences in aging generated by sex have long been recognized, but the underlying biological basis for these differences and the potential for sex-specific interventions remain understudied. To explore sex differences in the response to pro-longevity interventions, we utilized the *C. elegans* aging model and asked whether male lifespan and reproductive healthspan can be extended via compounds known to have pro-longevity effects in hermaphrodites. We tested seven different compounds at two concentrations each and found that lifespan was extended under all tested conditions. However, reproductive healthspan measured by mating success in late life improved under only two tested conditions, sulforaphane and metformin. These results demonstrate that lifespan and healthspan can be decoupled in *C. elegans* males and offer a new framework for screening pro-longevity compounds and for studying sex differences in aging in a classical aging model.

## Introduction

Sex differences in aging are robust and ubiquitous [1]. These differences are observed in both lifespan (the chronological time between birth and death) and healthspan (the proportion of lifespan that is disease-free). In humans, women live longer but tend to suffer worse health outcomes compared to men [2, 3]. Several age-associated conditions such as chronic conditions [4], bone mineral density loss [5], inflammation [6], and frailty [3] exhibit sex differences in incidence and severity [7]. This sexual dimorphism extends to modulators of aging such as pro-longevity interventions, with many tested regimes (genetic, chemical, or environmental) showing sex differences in efficacy [8]. However, our knowledge of the molecular basis of these differences is limited due to a historic focus on male health and disease [9].

*Caenorhabditis elegans* have a short lifespan (2-3 weeks), have conserved genetic pathways that modulate aging [10], have genetic diversity comparable to that of human populations [11, 12], and have longevity that is amenable to both genetic and chemical interventions [10], making them a pivotal model system for understanding the biology of aging. *C. elegans* is also sexually dimorphic, with self-fertile hermaphrodites and rare males [13, 14]. While a few recent studies have included sex differences in *C. elegans* aging [15–18], the vast majority of aging research using *C. elegans* focuses solely upon hermaphrodites. This is largely due to the fact that males are rare in the commonly used lab strain N2 (0.1%), as well as being shown to have detrimental effects on hermaphrodite lifespan [19–21]. Overall, our understanding of sex differences in aging processes in *C. elegans* remains extremely limited.

The *Caenorhabditis* Intervention Testing Program (CITP) is a National Institutes of Aging supported collaborative effort across three laboratories in the USA that leverages the benefits of *Caenorhabditis* nematodes as an aging model to test the effects of pharmacological interventions on the lifespan and healthspan of different species in the *Caenorhabditis* genus in a rigorous and robust fashion. The program has tested dozens of compounds and has found that many do indeed extend hermaphrodite lifespan [22]. However, it remains unknown whether these compounds are similarly efficacious in males. This seems especially pertinent because the CITP’s long-running sister program using mice, the ITP, has found that males tend to be more responsive to compound interventions than females [8, 23, 24].

Male *C. elegans* represent an exciting tool to test pharmacological interventions for two main reasons. First, males provide a complex health measure that integrates the health of multiple tissues and structures into one output: reproductive success. Male motivation for mating remains high throughout life and failure to sire progeny is caused by physical deterioration rather than a decrease in motivation per se [25, 26]. This is in contrast to hermaphrodites, which appear to change reproductive patterns due a variety of environmental inputs, including the presence of males [27]. Males need to locate a hermaphrodite via chemotaxis, scan their mate’s body to locate the vulva via mechanosensory neurons, insert the spicule, and ejaculate, thereby transferring sperm to the hermaphrodite uterus [28]. If any of these processes exhibit age-associated defects, reproductive success will decrease. The second reason males are an excellent tool for screening pharmacological compounds is that males do not produce self-progeny, eliminating the need to abrogate offspring production, for instance using the chemotherapy agent 5-Fluoro-2’-deoxyuridine (FUdR), which blocks cell division by inhibiting DNA synthesis [29, 30]. In hermaphrodite lifespan experiments, FUdR is commonly used to sterilize hermaphrodites and thereby eliminate the need for daily transfers during the first 5-6 days of adulthood (which is very labor and resource intensive). However, FUdR is known to affect hermaphrodites directly, as well as affecting the bacteria upon which they feed [31, 32]. Male lifespan experiments therefore offer the same ease as FUdR-supplemented hermaphrodites but without confounding effects of FUdR per se.

Here, we tested the potential pro-longevity effects of several CITP-validated compounds on male lifespan and healthspan, finding that all tested compounds increase male lifespan, even at lower concentrations than hermaphrodites. However, only two compounds reliably improved male healthspan, as measured via mating success in late life. The work here describes a new framework for pharmacological intervention screening using male *C. elegans* in addition to hermaphrodites to expand our understanding of sex differences in aging and to enable us to create interventions that are effective for both sexes.

## Materials and methods

### Worm strains and maintenance

All the strains used in this study were provided by the *Caenorhabditis* Genetics Center (CGC), which is funded by NIH Office of Research Infrastructure Programs (P40 OD010440). The strains used were CB1489 (*him-8(e1489)* IV) and JK574 (*fog-2(q71)* V). The strains were maintained on 60 mm plates of standard NGM media seeded with *E. coli* OP50-1. Worms were transferred to fresh plates 2-3 times weekly. Unless otherwise specified, all the stocks and assays were maintained at 20 ºC.

### Compound treatment

We used the following equation to determine the working stock concentration (X), where the treatment volume is 450 µL for water-soluble compounds and 25 µL for DMSO-soluble compounds and the plate volume is 10 mL for both types:

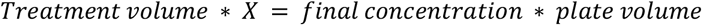

For water-soluble compounds (gold sodium thiomalate, metformin, thioflavin T), 450 µL of the working stock was added to 60 mM NGM plates seeded with OP50-1 and allowed to dry at room temperature for approximately 3 days. The working stock concentrations for metformin were 0.78 M and 1.56 M for final concentrations of 35 mM and 70 mM, respectively. The working stock concentrations for gold sodium thiomalate were 1.11 mM and 2.22 mM for final concentrations of 50 µM and 100 µM, respectively. The working stock concentrations for thioflavin T were 0.56 mM and 1.11 mM for final concentrations of 25 µM and 50 µM, respectively. For DMSO-soluble compounds (all *trans* retinoic acid, propyl gallate, resveratrol, sulforaphane), 25 µL of the working stock mixed with 425 µL of water was added to 60 mM NGM plates seeded with OP50-1 and allowed to dry at room temperature for approximately 3 days. The working stock concentrations for all *trans* retinoic acid for were 30 mM and 60 mM for final concentrations of 75 µM and 150 µM, respectively. The working stock concentrations for propyl gallate were 40 mM and 80 mM for final concentrations of 100 µM and 200 µM, respectively. The working stock concentrations for resveratrol were 20 mM and 40 mM for final concentrations of 50 µM and 100 µM, respectively. The working stock concentrations for sulforaphane were 20 mM and 40 mM for final concentrations of 50 µM and 100 µM, respectively.

### Lifespan assays

A modified CITP lifespan protocol was used [33]. Males at the L4 stage (identified by the distinct tail morphology in that stage) were selected from a mixed population plate and transferred to treated plates at a density of 40 worms per plate. Males were counted 3 times weekly, removing any dead worms, and transferred once weekly until all the worms were dead. Lost and walled worms and contaminated plates were censored from the final analyses.

### CITP data comparison

We calculated the change in survival and percent change in median lifespan in hermaphrodites using data obtained from the CITP Data Portal (citpaging.org) for each compound (and unpublished data for gold sodium thiomalate) and compared the results to the survival and percent change in median lifespan in males when treated with the same compound and concentration. We filtered the hermaphrodite data to include the *C. elegans* N2 strain data from the Oregon location only to minimize variability since the male lifespan experiments were conducted in Oregon. We additionally filtered the compound concentrations to include only those tested in males. The change in median lifespan was calculated from individual treatment replicates and their corresponding control replicates by subtracting the median lifespan of the control replicate from the median lifespan of the treated replicate then divided by the median lifespan of the control.

### Mating success assays

Males at the L4 stage (identified by the distinct tail morphology in that stage) were selected from a mixed population plate and transferred to treated plates at a density of 40 worms per plate 40. To generate seven-day old males, L4 males were moved to treated plates seven days prior to the assay. To generate five-day old males, L4 males were moved to treated plates five days prior to the assay. To generate one-day old males, L4 males were moved to treated plates one day prior to the assay. To generate virgin *fog-2* pseudo-females, L4 pseudo-females were moved to separate plates away from males one day prior to the assay to prevent them from mating. On the day of the assay, one male and two pseudo-females are placed on a 35 mm plate seeded with 100 µL of OP50-1 and allowed to mate. Approximately 24 hours later, the plates are scored for the presence or absence of progeny. For each compound, at least two trials (biological replicates) were conducted with approximately 20 plates (technical replicates) per trial.

### Statistical analysis

All the statistical analyses presented here were conducted using R version 4.4.1[34]. Lifespan data were analyzed using R scripts modified from the CITP’s standardized analysis pipeline [33]. To compare survival of each treatment group to the control group, a mixed effects Cox-Proportional Hazards model was used where the compound treatment was the fixed effect and both biological (Start Date) and technical (Rep) replicates were the random effects. To fit the mixed effects model, we used the coxme package version 2.2-20 [35]. Kaplan-Meier curve plots were generated using the survival package version 3.6-4 [36, 37]. To test for the effects of sex on the response to compound treatment, a mixed effects Cox-Proportional Hazards model was used where the compound treatment, sex, and the interaction were the fixed effects and both biological (Start Date) and technical (Rep) replicates were the random effects. To compare the mating success of each treatment group to the control group, a generalized linear mixed model with a binomial distribution was used where the compound treatment and age (and the interaction) were the fixed effects and biological replicate (Trial) was the random effect. To fit the linear model, we use the lme4 package version 1.1-35.5 [38]. All the scripts are available in Supplementary File 1.

## Results

### Hermaphrodite pro-longevity chemical compounds also extend lifespan in *C. elegans* males

We first asked if male lifespan can be extended via compounds known to have pro-longevity effects in hermaphrodites. Since the commonly used wildtype N2 strain has a low incidence of males (0.1%), we used *him-8* mutants to enrich for males in our populations. We selected seven compounds that have been previously shown by CITP and others to extend hermaphrodite lifespan. We tested the Vitamin A derivative all-*trans* retinoic acid [39], the anti-rheumatic compound gold sodium thiomalate (personal communication with CITP), the type 2 diabetes medication metformin [40], the antioxidants propyl gallate and resveratrol [33, 41], the cruciferous plant derivative sulforaphane [42, 43], and the amyloid dye thioflavin T [33, 44]. For each compound, we selected the lowest effective concentration previously reported to extend N2 hermaphrodite lifespan and asked whether those concentrations extend male lifespan in a similar manner. We found that all tested compounds significantly extended male median lifespan (Fig. 1, darker survival curves), with thioflavin T and sulforaphane resulting in the largest extension in median lifespan (76.92% and 43.75%, respectively) and maximum lifespan (100% and 42.86%, respectively). Notable extension in median and maximum lifespan was observed with all-*trans* retinoic acid (18.75% and 9.52%), gold sodium thiomalate (23.07% and 43.75%), metformin (23.07% and 43.75%), propyl gallate (18.75% and 14.29%), and resveratrol (25% and 33%) treatments. Since males have been shown to be more sensitive than hermaphrodites to certain chemicals [45, 46], we next asked whether a lower concentration of each compound was sufficient to extend lifespan. We lowered the concentration of each tested compound by half and found that lifespan was significantly extended at these concentrations as well (Fig. 1, lighter survival curves). Surprisingly, we found that the magnitude of median lifespan increases for gold sodium thiomalate, metformin, and thioflavin T was very similar at these lower concentrations, suggesting that males are more susceptible to chemical treatments and could represent a new tool for testing compound interventions in a more efficient and cost-effective manner.

**Fig. 1.**
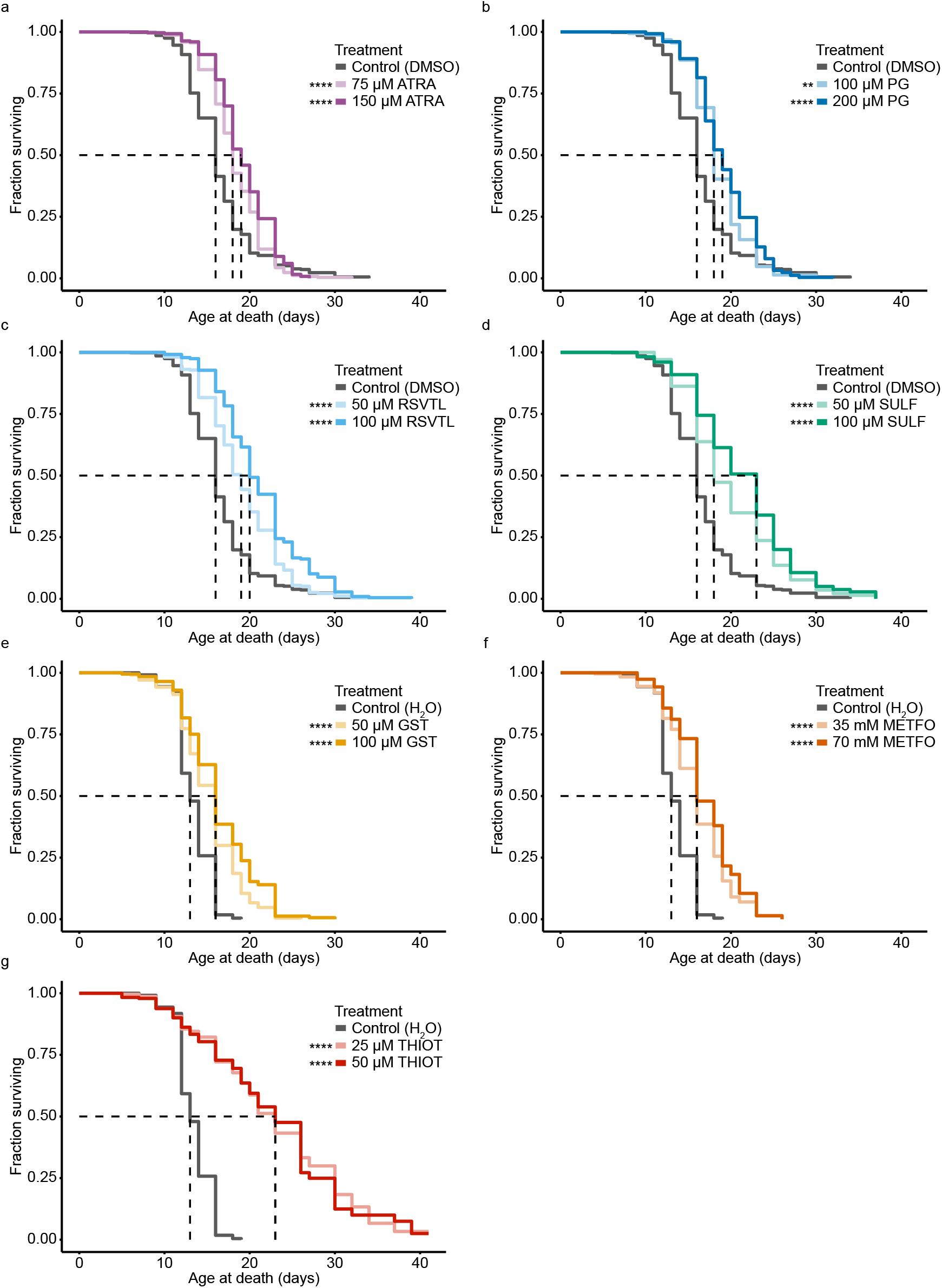
Pro-longevity compounds extend male lifespan. Kaplan-Meier curves showing survival of *C. elegans* males treated with either DMSO-soluble (a-d) or water-soluble (e-g) compounds. Solid lines represent survival on the negative control (DMSO or water), all-*trans* retinoic acid (ATRA), propyl gallate (PG), resveratrol (RSVTL), sulforaphane (SULF), gold sodium thiomalate (GST), metformin (METFO), and thioflavin T (THIOT). Each line represents at least two biological replicates with total n = 113-496. The dashed lines denote the age at which 50% of the population has died. The asterisks denote *p-*values from a Cox Proportional Hazards model where **p<.01, ****p<.0001. For additional information and the output of the CPH model, see Supplementary Table 1

While all the tested compounds extended male lifespan significantly, we wondered if the effect sizes for the compound treatment between males and hermaphrodites were different. Using previously published data for hermaphrodites from the CITP, we asked whether males respond quantitatively differently than hermaphrodites to compound treatments. We found that the magnitude of survival increase elicited by four compounds (all*-trans* retinoic acid, sulforaphane, gold sodium thiomalate, thioflavin T) is dependent on sex. Hermaphrodites treated with all*-trans* retinoic acid, sulforaphane, and gold sodium thiomalate display a larger increase in median lifespan than males. Males, on the other hand, display a larger increase in median lifespan than hermaphrodites when treated with thioflavin T (Fig. 2). Although the median lifespan change induced by metformin treatment is greater in hermaphrodites than males, greater replicate variance at the individual sampling level for metformin treatment reduced the power, therefore the difference itself was not significant. Overall, we find evidence for differences in the efficacy of compound treatment based on sex in *C. elegans*.

**Fig. 2.**
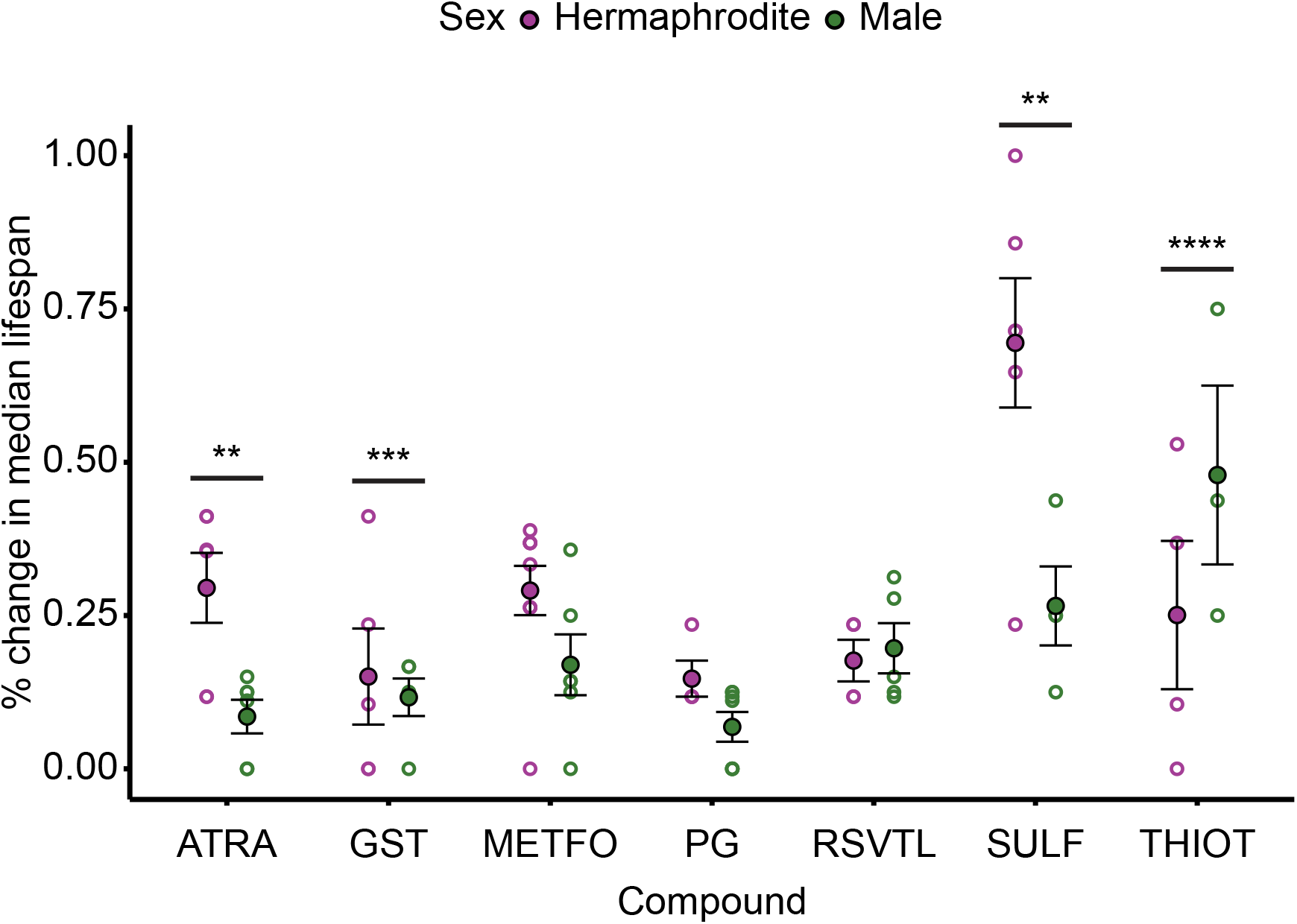
Sex differences in the percent change in median lifespan for different compounds compared to the control. Dot plot showing the percent change in median lifespan for each compound compared to the control. Solid circles represent the mean and open circles represent individual replicates. The error bars represent SEM. Only the higher concentrations (shown in Fig. 1) are represented here as lower concentrations were not tested on hermaphrodites. A CPH model was used to assess the effects of compound, sex, and the interaction of the two factors on survival. The asterisks denote *p-*values from the CPH model where **p<.01, ***p<.001, ****p<.0001. For additional information and the output of the linear model, see Supplementary Table 2

### Reproductive success is preserved late in life under sulforaphane and metformin treatment

We next asked whether the increase in lifespan is accompanied by an improvement in health, specifically reproductive health. We selected reproductive success in late life as our health measure as it is a complex behavior that declines precipitously with age [26]. We treated males with the seven selected chemical compounds and measured their reproductive success at three different ages: day 1 of adulthood when reproductive success is at its peak, day 5 of adulthood when reproductive success drastically declines, and day 7 of adulthood when reproductive success is severely diminished. Because hermaphrodites can produce self-progeny and do not depend on males for providing sperm, we used *fog-2* mutant hermaphrodites that fail to produce sperm, making them pseudo-females [47]. To measure reproductive success, a single male and two virgin *fog-2* pseudo-females were placed on a plate and allowed to mate for 24 hours. Plates were then screened for presence or absence of progeny, indicating successful mating or no/failed mating, respectively. We found that, despite the significant increase in lifespan with all treatments, only two of them improved reproductive success (Fig. 3). Sulforaphane increased mating success on day 7 by 201% percent and metformin increased mating success on day 5 by 215% percent (Fig. 3 d and f). Overall, we find that out of seven tested compounds that significantly increase male survival, only two improved male reproductive success in old age. This further validates the use of males to screen for pro-longevity interventions and the use of male mating success as a comprehensive healthspan metric.

**Fig. 3.**
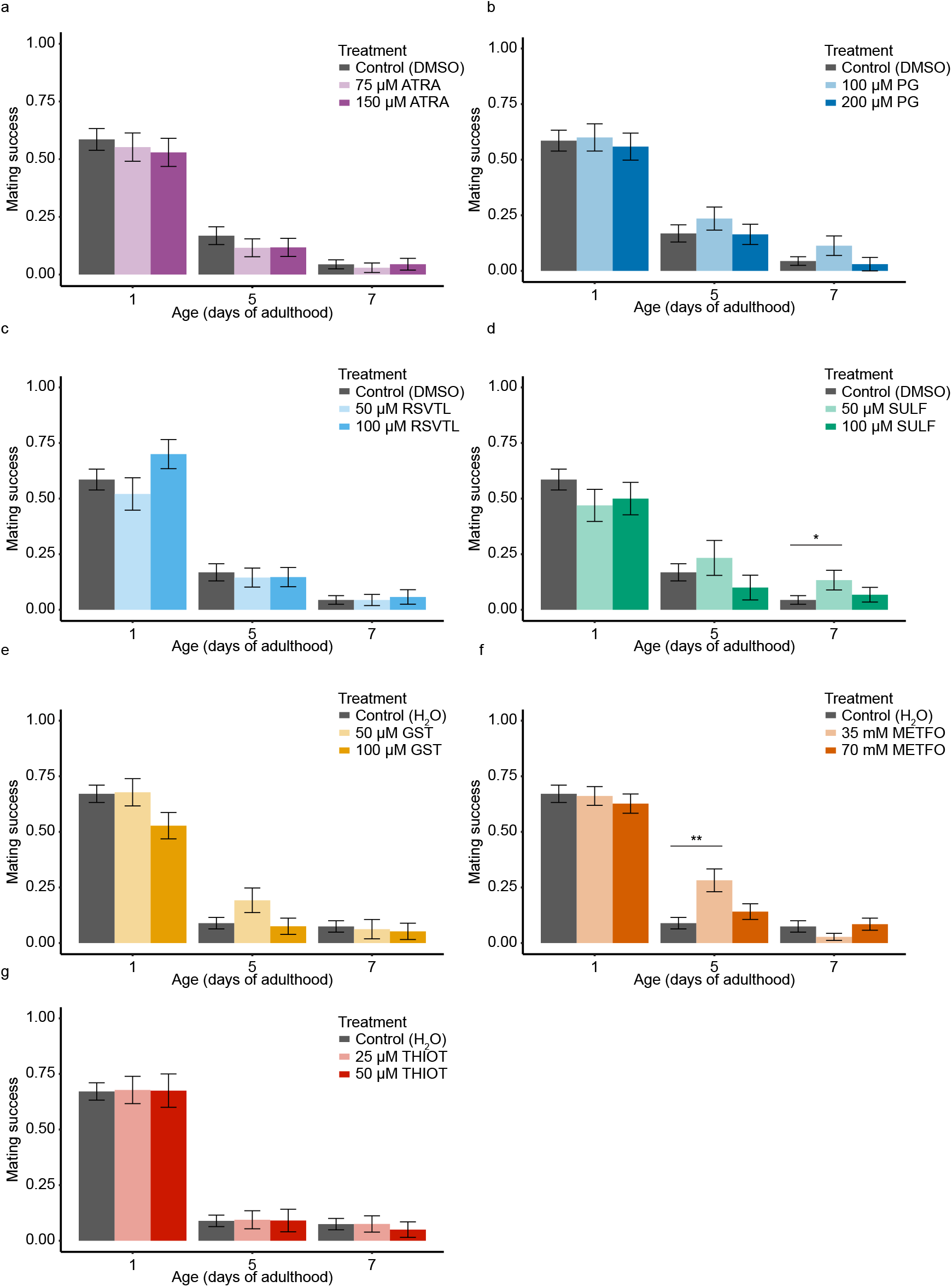
Sulforaphane and metformin preserve late-life male reproductive success. Bar graphs showing mating success for *C. elegans* males treated with either DMSO-soluble (a-d) or water-soluble (e-g) compounds then mated with *fog-2* pseudo-females. Each bar represents at least two biological replicates, with total n = 30-113. Error bars represent SEM. A generalized linear mixed model with a binomial distribution was used to assess the effect of treatment and age (and the interaction) on mating success. The asterisks denote *p-*values for age by compound interaction from the linear model where *p<.05, **p<.01. For additional information and the output of the linear model, see Supplementary Table 3

## Discussion

In this study, we leverage *C. elegans* males to investigate whether biological sex affects the response to a set of pro-longevity compounds validated by CITP. We found that treatment with all tested compounds in this study significantly extended male lifespan. Despite this, reproductive healthspan as measured by mating success was extended under only two tested compounds. To our knowledge, these findings represent the first effort to explore sex differences in response to pro-longevity compounds in the *C. elegans* model system.

Sex differences in biological processes have been historically neglected or ignored in both basic and clinical research [9]. This oversight was in part due to the incorrect assumption that female traits are generally more variable due to hormonal fluctuations [9]. Although this assumption was later debunked [48], the bias for using males in research persisted for many years, significantly limiting our knowledge of sex differences in biological processes and inappropriately generalizing male-specific findings to both sexes [49, 50]. To address this knowledge gap, the National Institutes of Health mandated in 1993 that women be included in clinical studies (National Institutes of Health 1994) and in 2015 that sex as a biological variable be included in study design, analysis, and reporting (National Institutes of Health 2015). Sex plays an especially important role in aging; there are robust differences between the sexes in lifespan, healthspan, age-associated disease susceptibility, and the response to pro-longevity interventions [7, 8]. Despite institutional efforts and scientific findings, research on sex differences in aging biology has remained fairly limited.

In 2002, the Intervention Testing Program (ITP) was created to screen compounds (chemical, pharmaceutical, or plant extracts) that extend lifespan or delay onset of diseases in a genetically heterogeneous mouse model [53]. The program emphasized robust and reproducible experiments and has identified 13 compounds that significantly increase median lifespan in mice [53, 54], many of which show sex differences in both the presence and magnitude of response. Of the ITP compounds that increase median lifespan, only five (rapamycin [55], meclizine [56], glycine [57], captopril [58], acarbose [23, 59, 60]) increase lifespan in both sexes, with rapamycin producing a greater response in females and captopril producing a greater response in males [54]. Several compounds, however, work exclusively in one sex; five compounds (NDGA [24, 60], Protandim [60], astaxanthin [56], Aspirin [24], 17-alpha-estradiol [23, 60, 61] extend median lifespan in males only, one compound ((R/S)-1,3-butanediol [58] extends median lifespan in females only, and two compounds (canagliflozin [62, 63], 16-alpha-hydroxyestradiol [63] extend male lifespan while reducing female lifespan [54]. Although the ITP successfully identified and validated the effects of many interventions, the need for a model for high-throughput screens emerged, leading to the formation of the CITP [22]. Inspired by the model of the ITP, the CITP exploits the genetic diversity of *Caenorhabditis* in addition to the short generation time and relatively low-cost of maintenance of nematode strains to conduct rigorous and reproducible experiments to identify compounds that extend lifespan and/ or healthspan on *Caenorhabditis* hermaphrodites. To date, the CITP has tested over 75 compounds and identified 12 that extend the median lifespan by at least 20% [22]. Of the 5 compounds tested by both the ITP and CITP, 3 significantly increased median lifespan in both species [22]. While the ITP tests compounds using both male and female mice, the CITP follows the convention of *C. elegans* work, using only hermaphrodites across multiple species for lifespan experiments. This has left a need to investigate sex differences in the response to compounds in this classical aging model. Here, we provide additional evidence for sexual dimorphism in the response to pro-longevity interventions. Of the seven tested compounds, sex modulates the effect size of four of them.

In the *C. elegans* aging field, males have been excluded from experiments due to multiple factors. First, males are rare in the commonly used N2 lab strain, representing around 0.1% of the population [14]. Second, Lifespan experiments can be complicated by the adverse effects of males on hermaphrodite lifespan, induced by mating or exposure to male pheromones [19–21]. Other males are also affected by male pheromones, leading to a density-dependent toxicity and lifespan decrease [46]. Third, males are highly motivated to search for mates, occasionally leading them to leave their agar media, crawl onto the plastic walls of petri dishes and desiccate. This leads to a decrease in the sample size, as these males are marked as “lost” and censored from final analyses. To circumvent the first problem, the male-enriched *him-8* strain was used in this study. The HIM-8 protein binds the meiotic pairing center of the X chromosome, and its loss leads to an increase in X chromosome nondisjunction events and a higher incidence of males [64]. Although we note the negative effect of male-male interactions on lifespan, this was not severe enough to abolish the positive lifespan extension effects of the compounds tested in this study and actually highlights the efficacy of these interventions. Future work could investigate the effects of these compounds on males with mutations in *daf-22* that are deficient in pheromone production, thereby causing weaker male-induced demise [20, 46]. Finally, in our hands on average 10% of males were marked “lost” (excluding losses due to contamination) and censored from the lifespan experiments, which is not high enough to interfere with robust assessment of the treatments.

Despite these potential challenges, males provide a unique tool for measuring the effects of compounds on lifespan and healthspan. One of these benefits is eliminating the need for progeny abrogation using FUdR thus eliminating its confounding effects on lifespan because males, unlike hermaphrodites, are not capable of producing self-progeny [31, 32]. Males additionally provide a unique measure for reproductive healthspan: mating success. Mating is a complex behavior that is controlled by multiple tissue types and motivated by a consistent drive. Indeed, males lose their ability to mate in old age due to neuromuscular deficits and not due to a decrease in drive or sperm quality [25, 26]. Mating ability offers a complex yet easily-screenable phenotype (presence or absence of progeny) that could indicate a systemic healthspan extension. We show here that reproductive success in late life can be modulated by compound interventions, thus validating male mating success as a viable healthspan measure for future studies. Given the sensitivity of the system and the number of biological processes that modulate this phenotype, it could potentially be used as a highly sensitive screening tool for healthspan effects independent of lifespan extension.

The importance of describing both lifespan and healthspan while measuring the effects of pro-longevity interventions has been gaining increasing attention within the aging field [65–68]. Our findings further highlight the independent nature of lifespan and healthspan, as our study provides further evidence that lifespan extensions are not necessarily coupled with healthspan extensions. In fact, the compounds tested in this study mostly increased lifespan without improving the healthspan measure used here. Because our reproductive healthspan measure encompasses a number of systems, such as neurological and muscular functions, that are of central interest in human aging [69, 70], this decoupling offers a potential opportunity. Indeed, it should be possible to screen for compounds that enhance late-life male reproductive function more directly, which could reveal novel pathways for future interventions. Future work could investigate the molecular mechanisms and behavioral changes underlying the improved reproductive success in late life induced by sulforaphane and metformin. Additionally, future work could investigate whether these compounds improve any other health metrics such as motility, oxidative stress resistance, and other standard healthspan measures [65].

In conclusion, we describe here the first study to use male *C. elegans* to investigate the effects of pro-longevity compounds on lifespan and reproductive healthspan. We emphasize the benefits, which largely outweigh the challenges, of including males in future studies and analyses especially when asking questions about aging. This will add to the fundamental, yet limited, knowledge of how sex influences biological processes in the classical *C. elegans* aging model. Ultimately, this will bring us one step closer to identifying interventions and regimes that benefit both sexes.

## Acknowledgements

We thank the CITP members Christine Sedore and Anna Coleman-Hulbert at the University of Oregon for their support and guidance throughout this project, their advice on data analysis, and their help preparing compounds and media. Some data were downloaded from the CITP Data Portal (citpaging.org), which is supported by funding from NIH’s NIA Division of Aging Biology (U24AG056052). Strains were provided by the CGC, which is funded by NIH Office of Research Infrastructure Programs (P40 OD010440).

